# High protein diets early in life improve survival, reduce cannibalism, and increase growth and yield in farmed yellow mealworms

**DOI:** 10.1101/2025.11.09.687235

**Authors:** N Dobrowolski, CD Perl, J Brumfield, J Christie, R Rodriguez-Guevara, SO Durosaro, M Barrett

**Affiliations:** Department of Biology, Indiana University Indianapolis, Indiana, USA; Institute of Aquaculture, University of Stirling, Stirling, UK; Department of Biology, Ivy Tech Community College, Indiana, USA

**Keywords:** insect welfare, yellow mealworms, nutrition, juvenile development, cannibalistic behavior

## Abstract

In the wild, animals may be able to compensate for nutritional inadequacies through foraging for novel resources; however, on farms, animals have more limited nutritional options. Therefore, providing adequate and accessible nutrition is essential to positive farmed animal welfare. Billions of yellow mealworms (*Tenebrio molitor;* Coleoptera: Tenebrionidae) are farmed as mini-livestock each year for use as food and feed. The major feedstock used, wheat bran, can sustain complete development of the species, but has been shown to have significantly lower protein content than yellow mealworms would typically self-select, particularly at early developmental stages. This can reduce growth, survival, and adult oviposition, and may contribute to high rates of cannibalistic behaviour among larvae (as larvae may have few options but to seek protein from the bodies of conspecifics). In this study, we use 20% nutritional yeast as a dietary protein supplement for larvae in early (0-4 weeks) and later life (4-8 weeks) to assess the effects on survival, growth rate, and rates of cannibalism, as well as any effects on growth, development timing, and adult body size. We also investigated the impact of this higher protein diet, when fed to either larvae or adults, on fecundity. We find that protein supplementation in larval diets increased survival by 23%, growth rate by 200%, and total biomass at eight weeks by up to 272%. Larval cannibalistic behaviour was reduced and development time was shorter. The high protein diet, whether fed to adults or to larvae, increased the number of offspring per female by 34-36%. Survival and growth benefits of protein supplementation were especially pronounced when fed to early juvenile instars, but can benefit mealworms across the entire first eight weeks of development. We highlight the need for further research into dietary protein supplementation for yellow mealworms across development to better align with their dietary choices, thereby improving fitness, yield, and welfare.

## Introduction

Yellow mealworm (*Tenebrio molitor;* Coleoptera: Tenebrionidae) larvae (YML) are one of the most abundantly farmed insects, with an estimated 290 billion reared annually as of 2020 (Rowe, 2020). This number is predicted to grow, with YML potentially playing a significant role in human, chicken, or pig food chains (Hong *et al*., 2020) as well as a fishmeal replacement (Shafique *et al*., 2021). While insect sentience is still debated (Eisemann *et al*., 1984; Adamo, 2016; Gibbons *et al*., 2022; Crump *et al*., 2023; Barrett & Fischer, 2025; Fischer *et al*., 2025), especially at the larval stage, many producers and academics advocate for a precautionary approach to insect welfare in the industry (IPIFF, 2019; van Huis, 2021; Barrett & Fischer, 2023). Growing evidence indicates that consideration of insect welfare can also be important for producers economically via productivity gains or social and regulatory impacts, given public interest in insect welfare (Barrett & Adcock, 2023; Niyonsaba *et al*., 2024).

One of the most widely employed frameworks for assessing animal welfare, the five domains (Mellor & Reid, 1994), places adequate nutrition (both quantity and quality) as a central component of high welfare. In industrial conditions, YML are frequently provided with wheat bran (Deruytter *et al*., 2021), a common byproduct of flour production (Prückler *et al*., 2014). The use of agricultural byproducts is key to the industry’s sustainability goals (DEFRA, 2023; Smetana *et al*., 2023; Hancz *et al*., 2024), however, these may not always provide adequate nutrition for developing larvae (Barrett *et al*., 2023). This can have significant impacts on larval growth, survival and nutrient content (Tamim *et al*., 2025). Wheat bran contains all the required nutrients for YML growth but not in sufficient ratios (Morales-Ramos *et al*., 2010; Ribeiro *et al*., 2018), particularly being too low in protein. YML had improved growth rates, food conversion efficiency, and fecundity when diets were at least 20% protein (Morales-Ramos *et al*., 2010), while wheat bran offers only 13-18% protein (Prückler *et al*., 2014).

Insects in the wild may resolve imbalances in ideal nutrient ratios behaviourally (e.g., self-selection, Simpson *et al*., 1988; Lee *et al*., 2012), and there is evidence that adult *T. molitor* can do the same (Rho & Lee, 2016). In general, the ability of individual insects to compensate for nutritionally-deficient diets has been well-explored via geometric nutritional frameworks (Raubenheimer & Simpson, 2003; Behmer, 2009; Simpson *et al*., 2015). Choice experiments show that insects can ingest mixtures of suboptimal diets, if such a mixture is available, to ensure consumption of their preferred macronutrient ratios (Behmer, 2009). This presents a problem in industrial rearing environments, where individuals have fewer options for varying their dietary intake. In fact, the sole way that YML may be able to adjust their nutrient intake, particularly under starvation conditions (Yang *et al*., 2018), is to resort to cannibalistic behaviour (Weaver & McFarlane, 1990; Randall *et al*., 2023).

Cannibalism of live individuals creates cuticular openings which are easily exploited by infectious agents, creating a significant disease risk (Williams, 2008; Barrett *et al*., 2023). Cuticular openings can cause death via haemolymph loss or dehydration over several days (Ichikawa & Kurauchi, 2009). Internal wounds from cannibalism could also cause a slow death via septicemia (Weaver & McFarlane, 1990). Cannibalism may also spread disease to healthy, live individuals when they consume dead or decaying conspecifics (reviewed in Barrett *et al*., 2023). Beyond these welfare impacts, low-protein rearing conditions may also have a negative effect on production if they reduce growth (van Broekhoven *et al*., 2015), survival (potentially reducing total biomass), or increase development time which may delay reproduction, slaughter, or sale (Morales-Ramos *et al*., 2010).

There is evidence that providing supplementary protein can improve growth rates and increase larval survival (Morales-Ramos *et al*., 2013; Oonincx *et al*., 2015; van Broekhoven *et* al., 2015; Rho & Lee, 2022), however, there are scant details available regarding the extent to which protein-supplemented diets are employed in industrial settings (Barrett *et al*., 2023). Where data are reported, it is unknown to what extent additional protein must be provided throughout development. Responses to high-protein diets during early compared with late juvenile developmental stages differ across insect herbivores (Woods, 1998). For instance, in Lepidopteran larvae, access to protein during early development may be more crucial for growth and survival than access during later instars; further, individuals may acclimate to either high or low protein, and therefore switching diets during development may prove detrimental (Woods, 1998). Dietary preferences may also shift during development: yellow mealworms show a preference for higher protein ratios (1:1 P:C) at early instars (compared to 1:1.24 at later instars; Rho & Lee, 2022), a preference matched by pine beetle larvae (Zhang *et al*., 2025) and *Lymantria* moth larvae (Stockhoff, 1993). Thus the timing of protein supplementation during YML development, as well as any benefits of that supplementation, remain an open question.

To address the potential benefits of protein supplementation for YML, we assessed the impacts of supplementing the usual wheat bran diet with 20% yeast, a commonly reported laboratory protein supplement (Ribeiro *et al*., 2018). In larger bins, we assessed the effect of supplemental yeast on larval growth rate, development time, adult body mass, and adult fecundity. We also assessed if early life (weeks 0 - 4) or later life (weeks 4 - 8) protein supplementation had a larger impact on evidence of cannibalistic behaviour, larval growth rate, survival, and total biomass. Finally, we assessed the impact on fecundity of protein supplementation in the diets for adults that did not receive juvenile protein supplementation.

## Methods

### Larval growth and development over 11 weeks

Animals were reared at room temperature and humidity (22.7 ± 0.4 °C and 63.9 ± 6.3% RH), under natural lighting cycles supplemented with room light (which amounted to a 14:10 L:D cycle during the experimental period).

Yellow mealworm eggs were sourced from Fluker Farms with an approximate hatch date. One day before the hatch date, approximately 3.5 g of eggs were placed on 100 g of feed in 25 oz plastic containers; the feed was either 100% wheat bran (low protein; LP diet) or 20% nutritional yeast (Bulk Priced Food Shoppe) and 80% wheat bran (Baker’s Bran) by weight (high protein; HP diet). Larvae were provided with 30-35 g of polyacrylamide gel three times a week for hydration. Polyacrylamide gel crystals (Oycevila) was used instead of agar gel as it has less protein; pilot tests with our lab population did not indicate impacts on mortality or growth (and this gel is frequently provided for rearing insects, e.g., Fluker Farms ‘Cricket Quencher’).

At 4 weeks, larvae were provided with 200 g of fresh feed and moved to 1.5 gallon rubbermaid plastic containers. Another 200 g of new feed was added at seven weeks. At 9 weeks, larvae were sifted from the old feed and frass and returned to 300 g of fresh feed; this was repeated weekly thereafter. 75-80 g of polyacrylamide gel was added four times a week starting in week 4.

Weekly, starting in week 2, three samples of ten larvae were taken (from the left, center, and right of the bin following some by-hand mixing of the substrate) from each bin and weighed to obtain the average larval weight. Weights were collected until the bins all reached ‘slaughter weight’ which we defined as 100 mg/larvae (Zacarias et al., 2025). Although we attempted to measure total biomass and estimated survival at week 11, polyacrylamide gel fragments in the feed prevented efficient sieving and total biomass data were considered too unreliable to be used for either purpose. Data were recorded twice in 24 hours from each bin during week 11 due to suspected non-random sampling during that week. We present both sets of week 11 data points (Figure 1), with the suspected-errored points in grey for transparency; however, only the second data points were analysed statistically.

**Figure 1.**
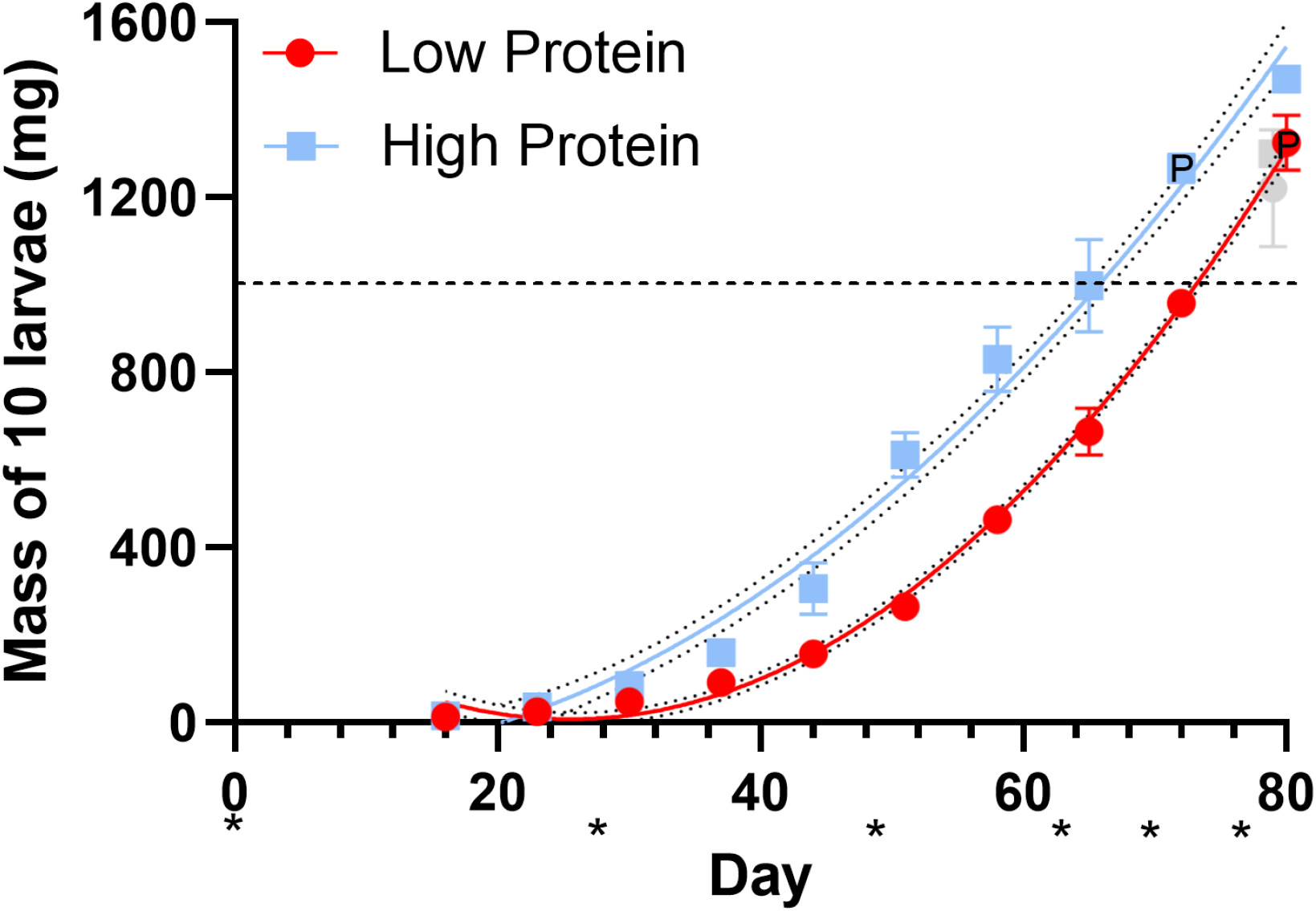
Larvae fed the HP diet reach slaughter weight faster than larvae fed the LP diet. Larvae fed the HP diet (blue) reached a typical slaughter weight after 66 days compared to 73 days on the LP diet (red; dotted lines are 95% CI). Growth rate for the first 11 weeks was faster on the high protein diet than the low protein diet (p < 0.0001). Asterisks indicate fresh feed was provided; the black dashed line indicates slaughter weight (100 mg/larvae). P indicates pupae were present in the sample when that datapoint was collected, however only larvae were weighed.

### Adult body mass and development time

To obtain development time and information on adult mass, bins from the prior study were monitored daily for adult emergence. Each day, adults were removed from the substrate and the total number that emerged was recorded; the first 100 adults from each replicate were weighed and sexed. All remaining adults were counted and weighed together as a batch so average adult weight for the day could be obtained. When a bin had fewer than 5 adults emerge per day for 2 consecutive days, that bin was removed from the study.

### Effects of early- vs. late-development diet on larval growth rate, survival, and cannibalism (weeks 4 - 8)

Animals were reared at room temperature and humidity (22.7 ± 0.4 °C and 64.6 ± 5.9% RH), under natural lighting cycles supplemented with room light (which amounted to a 14:10 L:D cycle during the experimental period).

200 larvae were removed from a large bin at four weeks of age (their early life diet followed description above in ‘Larval growth and development over 11 weeks’) and separated into groups of 10. Each group of ten was weighed and randomly assigned to receive either a LP diet or HP diet for the next four weeks, such that 100 larvae were assigned to each treatment.

This process was repeated for each of the large bins, generating treatments where larvae were assigned to a diet that either matched or differed from their early-development diet. This generated a 2x2 matrix (LP-LP, LP-HP, HP-HP, HP-LP) with five replicates per condition (one replicate of each type per large bin replicate from the above experiment).

Late-development (e.g., weeks 4 - 8) LP diets received 10 g of wheat bran and 2-3 g of polyacrylamide gel in a 90 mm diameter petri dish with the lid vented for air flow. HP current diets received 8 g of wheat bran, 2 g of nutritional yeast, and 2-3 g of polyacrylamide gel. 90 mm petri dishes were kept in secondary, 160 mm diameter petri dish lids with the sides coated with Fluon (byFormica, FluonPlus) to prevent escape (though larvae were rarely found outside their substrate in the 90 mm dishes). Gel was replenished every 4 days for the first two weeks and every two days for the last two weeks.

Every 48 hours, the number of live and dead larvae in each petri dish was individually counted. Live larvae were weighed as a group and returned to the feed. Dead larvae were examined for evidence of cannibalism (e.g. missing sections of body parts) and subsequently excluded from the rearing petri dishes. If any larvae could not be found live or dead, the condition was counted a second time. The number of missing larvae was recorded; if that larva was never found again for the duration of the experiment, it was logged as permanently ‘missing’. For analyses where we assessed the ‘minimum’ possible dead and/or cannibalized, the missing were not included. For analyses where we assessed the ‘maximum’ possible dead and/or cannibalized, the missing were included.

Old feed was removed and new feed provided at two weeks and three weeks, with the experiment terminating after four weeks (8 weeks of age). Whenever new feed was provided, the original counter counted the individuals twice before transfer and an additional researcher (MB) sorted through the old feed twice more to ensure there were no remaining live or dead larvae.

### Effect of larval and adult dietary supplementation with yeast on oviposition

Larval/pupal animals were reared at room temperature and humidity (22.7 ± 0.5 °C and 64.9 ± 6.0% RH), under natural lighting cycles supplemented with room light (which amounted to a 14:10 L:D cycle during the experimental period); adult animals for the experiment testing the effects of the larval diet were reared at room temperature and humidity (22.5 ± 1.4 °C and 68.2 ± 2.6% RH) and the same for the second experiment testing the effects of adult diet (22.5 ± 1.6 °C and 68.4 ± 3.0% RH).

Replicates of larvae reared on HP and LP diets were assigned to this experiment after they began to produce at least 30 adults a day for a three day period; then, fecundity was assessed as in Mahmoud *et al*., (2025). Briefly, 30 adults that were 0-24 hours old were placed into a plastic rearing box (14 L x 8.6 W x 4.8 H cm). They were provided with 35 g of either wheat flour (LP) or wheat flour and 20% yeast (HP) and a small mesh excluder (1 mm) through which females could lay eggs in order to avoid cannibalism of the eggs by adults. Moisture was provided via *ad libitum* polyacrylamide gel atop a small piece of plastic wrap.

In the first experiment testing the effects of larval diet, adults from the HP or LP larval replicates were provided with a LP diet after eclosion. They were allowed to oviposit for 15 days, after which point all adults were removed, sexed, and the substrate was sieved for eggs and larvae which were frozen and then counted under a microscope. From each rearing replicate, three adult replicates (n = 90 adults) were made, resulting in an n = 15 boxes/larval diet. One replicate (LP1-B) was removed from the analyses as a large number of individuals could not be accurately sexed from that condition.

For the second experiment assessing the effect of adult diet, all adults were sourced from LP larval replicates. Adults were provided with either a LP or HP diet after eclosion. Egg counting proved challenging in the previous experiment due to hatching of eggs into larvae, therefore oviposition time was limited to 10 days for this experiment. Additionally, we covered half the container with black Cinefoil to provide a shelter. After 10 days, all adults were removed, sexed, and the diet was sieved for eggs and larvae which were counted. From each rearing replicate, three adult replicates (n = 90 adults) were made, resulting in an n = 15 boxes/adult diet.

The number of offspring per female was determined by dividing the total number of unhatched eggs and larvae by the number of adult females in the box.

### Statistical Analyses

All statistics were run in R 4.4.2 or GraphPad Prism v. 10.4.2. Alpha was set to 0.05 for all analyses. Data were evaluated for normality and heteroskedasticity when determining which tests to run.

For growth rate across the first eleven weeks, an extra sum-of-squares F-test was used to assess if there was a difference in the line of best fit for HP v. LP-fed larvae and if a non-linear or linear regression was the best fit to model growth under each diet. An unpaired t-test was used to assess differences in mass at 11 weeks between diets.

For growth rate from weeks 4 - 8, an extra sum-of-squares F-test was used, without including intercept, to assess if there was variance among conditions in growth rate (best-fit second-order polynomial lines) or if a single line could be used to model all datasets. Average growth per day and total biomass were analyzed across conditions using an ANOVA followed by a Tukey’s post hoc Multiple Comparisons test. Survival was measured as the proportion of larvae remaining alive on the last day of the experiment. Proportion surviving and proportion cannibalised were analysed using a beta regression (Cribari-Neto & Zeileis, 2010), with model term significance ascertained using a type II ANOVA (Fox & Weisberg, 2019). Non-significant terms were eliminated piecewise from the model to yield a minimum adequate model. Post-hoc pairwise comparisons were conducted using the function ‘emmeans’ from the package “emmeans” (Lenth, 2024).

Differences in the number of adults produced per bin, and the mean eclosion day (development time) per bin, were assessed using an unpaired t-test. For the effect of diet on adult body mass over time, the data presented non-normally distributed residuals when analysed using a linear model (ANCOVA); however, we found that there was no significant difference between using a linear model or a generalised linear model, in either results or explanatory power. Therefore, results from the simpler linear model (ANCOVA) are presented here.

Numbers of eggs laid per female were analysed using linear mixed effects models (Bates *et al*., 2015; Kuznetsova *et al*., 2017), with larval replicate treated as a random effect. Post-hoc pairwise comparisons were conducted using the function ‘emmeans’ from the package “emmeans” (Lenth, 2024).

## Results

### Larval growth rate over 11 weeks

Growth rate for the first 11 weeks was faster on the high protein diet than the low protein diet (F_94,100_ = 109.2, p < 0.0001). The best-fit line for LP growth was: y = 0.44 x^2^ - 22.24 x + 290.5 (F_47,50_ = 1018, p < 0.0001; R^2^ = 0.994) and predicted larvae reaching slaughter weight (100 mg) on day 73; the best-fit line for HP growth was: y = 0.27 x^2^ - 1.315 x - 84.68 (F_47,50_ = 87.03, p < 0.0001; R^2^ = 0.98) and predicted larvae reaching slaughter weight on day 66 (Figure 1). Larvae reared on the LP diet were smaller at 11 weeks, with a body mass of 132.5 ± 6.37 mg compared to 147 ± 2.56 mg for the HP diet (unpaired t-test: t = 4.7_10,8_, p = 0.0015); HP larvae had an 11% increase in mass.

### Growth rate, survival and cannibalistic behavior from weeks 4 - 8

Mass over time varied among the four conditions (LP - LP, LP - HP, HP - HP, and HP - LP; Extra sum of squares F-test: F_286, 298_ = 80.16, p < 0.0001). A second order polynomial was a better fit than a straight line (all F_71,74/75_ > 65.72, p<0.0001; Figure 2). Average growth rate (mg/day over the 4 week time period; Figure 2) was highest in the HP-HP condition, followed by the HP-LP condition, LP-HP condition, and the LP-LP condition (ANOVA, F_16,20_ = 48.75, p < 0.0001; Tukey’s MCT: all p < 0.05; Figure 2). Growth per day was doubled in the HP-HP condition (2.01 ± 0.2 mg/larva/day) compared to the LP-LP condition (Table 1; 1.00 ± 0.1 mg/larva/day).

**Table 1.** Survival, growth per larva, and total biomass in each condition from weeks 4-8.

| Condition | Survival (%; mean +/- SD) | Growth (mg/larva/day) | Total biomass (g) |
| --- | --- | --- | --- |
| LP - LP | 69.60 ± 12.82 | 1.00 ± 0.10 | 2.44 ± 0.59 |
| LP - HP | 82.05 ± 6.62 | 1.38 ± 0.10 | 3.85 ± 0.16 |
| HP - HP | 93.02 ± 3.47 | 2.01 ± 0.20 | 6.63 ± 0.66 |
| HP - LP | 88.40 ± 3.98 | 1.67 ± 0.14 | 5.31 ± 0.38 |

**Figure 2.**
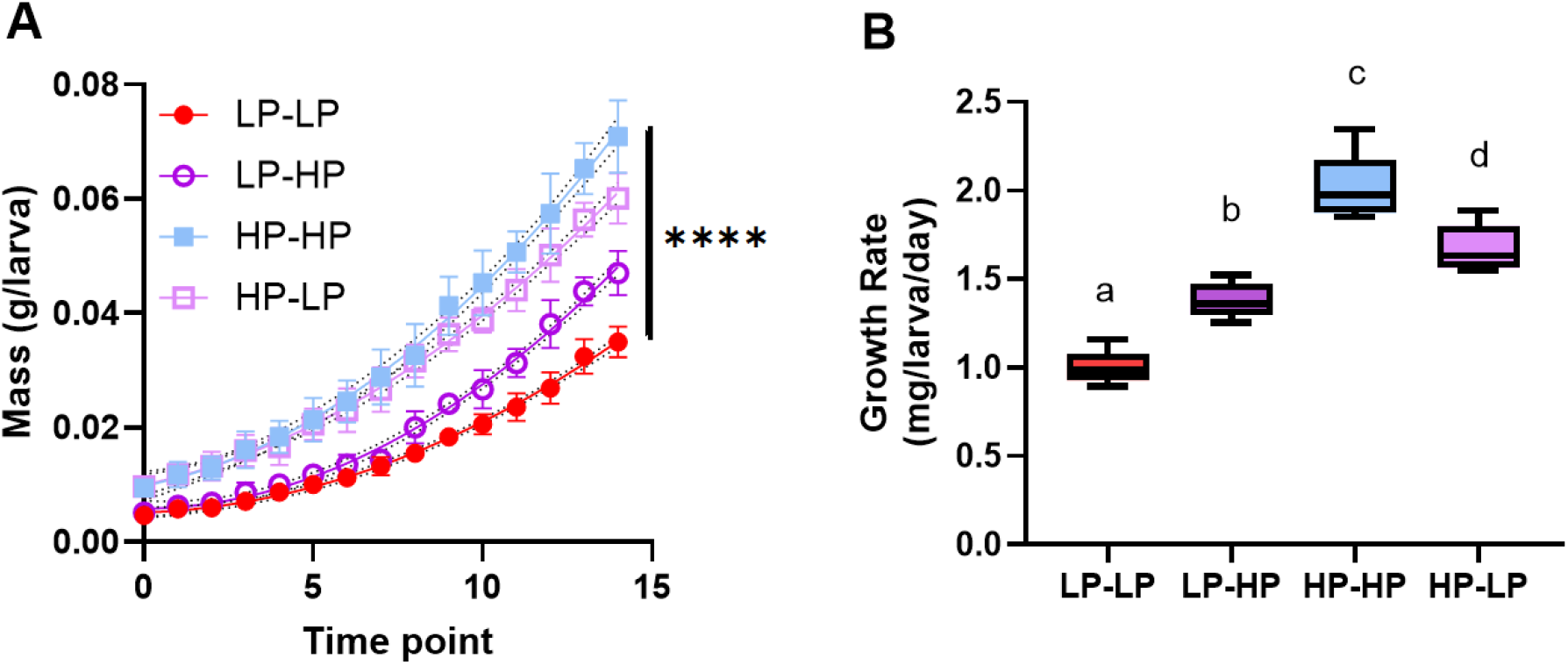
Early life and current diet affect the growth rate of YML. **A)** Overall models of growth rate in each condition differed based on both early life diet and current diet (Extra sum of squares F-test: F_286, 298_ = 80.16, p < 0.0001). **B)** Average growth per larva per day was highest in the HP-HP treatment, then the HP-LP treatment, then LP-HP, and finally LP-LP. Letters indicate statistically significant differences (ANOVA, F_16,20_ = 48.75, p < 0.0001; Tukey’s MCT: all p < 0.05).

Both early life diet (beta regression: z = -5.04, p < 0.0001) and current diet (beta regression: z = -2.93, p = 0.003) significantly affected survival from week 4 to week 8, but not their interaction (which was dropped from the model; beta regression: z = -0.36, p = 0.72). LP diets early in life reduced survival (beta regression: z = 5.26, p < 0.0001; main effect, averaged across levels of current diet), as did LP diets in weeks 4 to 8 (Figure 3A; z = 2.97, p = 0.003; main effect, averaged across levels of early life diet).

**Figure 3.**
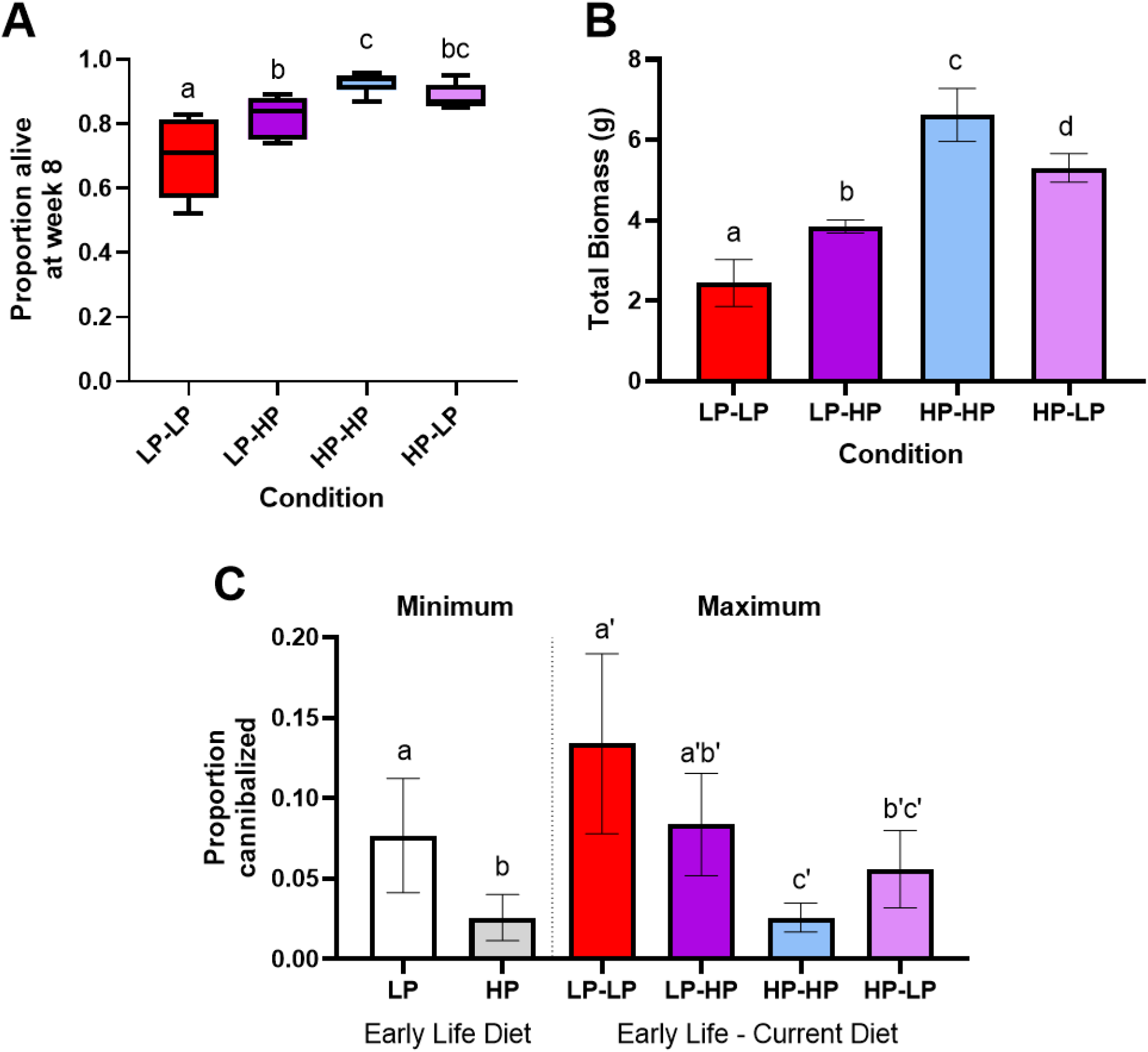
Effects of early life, and current, diet on survival, total biomass, and cannibalistic behaviors in yellow mealworms. A) Low protein diets early in life, and during weeks 4 - 8, reduced survival in yellow mealworms (beta regression with Tukey’s MCT; all p < 0.05). B) Total biomass produced varied based on both early life and current diet (ANOVA with Tukey’s MCT; all p < 0.01). C) Early life diets increase the likelihood of cannibalism. The minimum possible proportion of larvae cannibalized (direct evidence of cannibalism only) was decreased by HP diets early in life. The maximum possible proportion of larvae cannibalized (direct and indirect evidence of cannibalism) was decreased by HP diets currently and early in life (beta regression with Tukey’s MCT, all p < 0.01). A-C) Letters indicated statistically significant differences (p < 0.05) among conditions on a graph.

Variation in survival and growth resulted in large differences in total biomass produced by week 8 across all conditions (Figure 3B; Table 1; ANOVA: F_16,20_ = 69.91, p < 0.0001), with HP diets for 8 weeks having 2.7-fold biomass production compared to LP diets for 8 weeks (Tukey’s MCT: q = 19.33, p < 0.0001).

The minimum proportion of cannibalized larvae was significantly higher for the early life diet treatment than the HP early life diet treatment (Figure 3C; beta regression: z = 4.64, p < 0.0001). The best-fit model for the maximum proportion (observed cannibalized dead larvae and missing larvae) of larvae cannibalized was increased. The maximum proportion of cannibalized larvae was significantly higher in treatments with LP early life diet (beta regression: z = 5.18, p <0.0001) and LP current diet (beta regression: z = 3.11, p = 0.0019).

### Number of adults, development time, adult body mass

There was no difference in the number of adults produced in the large HP and LP bins (unpaired t-test; t_8,10_ = 1.65, p = 0.14); each bin produced a mean of 1602 ± 134 beetles (mean ± SD).

Mean development time per bin was faster in the HP conditions compared to the LP conditions (Figure 4; unpaired t-test; t_8,10_ = 36.09, p < 0.0001). Adults in the high protein condition emerged on a mean of 23.31 ± 0.72 days (mean ± SD) following the emergence of the first beetle in any condition; adults in the low protein condition emerged on a mean of 36.46 ± 0.39 days.

**Figure 4.**
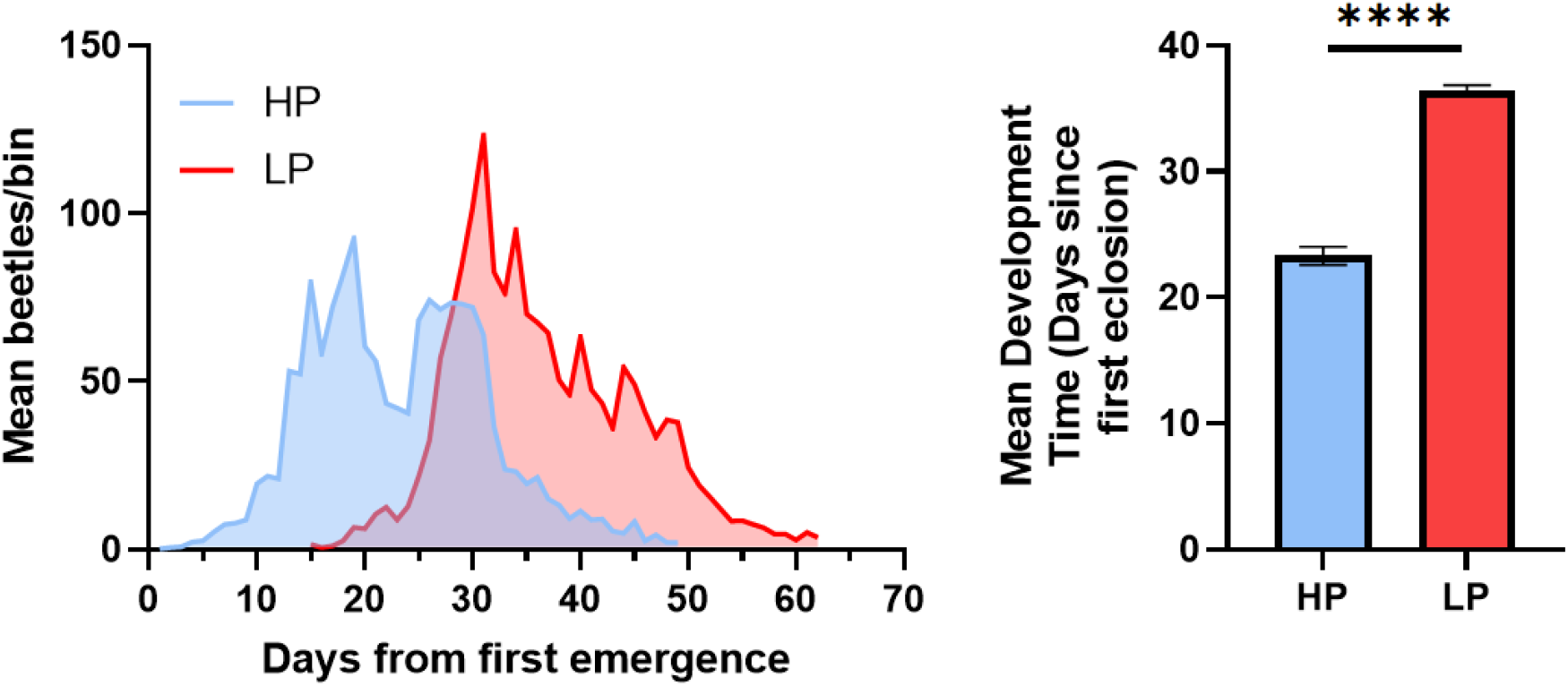
Mean eclosion day is 13 days faster in HP fed mealworms than LP fed mealworms. Adult mealworms from the HP fed condition emerged at 23.31 ± 0.72 (mean ± SD) days following the emergence of the first beetle in any condition; adults in the low protein condition emerged on a mean of 36.46 ± 0.39 days (unpaired t-test; t_8,10_ = 36.09, p < 0.0001). **** = p < 0.0001

The observation of two ‘peaks’ in the high protein condition, as well as smaller adults in the first ‘peak’ period, suggests that feed may not have been *ad libitum* in the high protein conditions despite being given the same mass of feed per larvae as the low protein conditions, perhaps due to a faster growth rate of the HP-fed larvae.

For each condition, there was a significant increase in the mean adult mass per day over time (ANCOVA; t_445,449_ = 25.09, p < 0.0001). There was a significant difference in the mean mass on emergence among treatments (ANCOVA; t_445,449_ = 2.67, p < 0.001). Mean emergence mass per day of adults from the LP treatment (mean ± SD = 0.11 ± 0.013 g) was higher than for the HP treatment (0.10 ± 0.015). There was a significant interaction between diet and emergence date (ANCOVA; t_445,449_ = 2.68, p < 0.001). The increase in mean adult emergence mass per day was steeper in the high protein treatment, than the low protein treatment.

**Figure 5.**
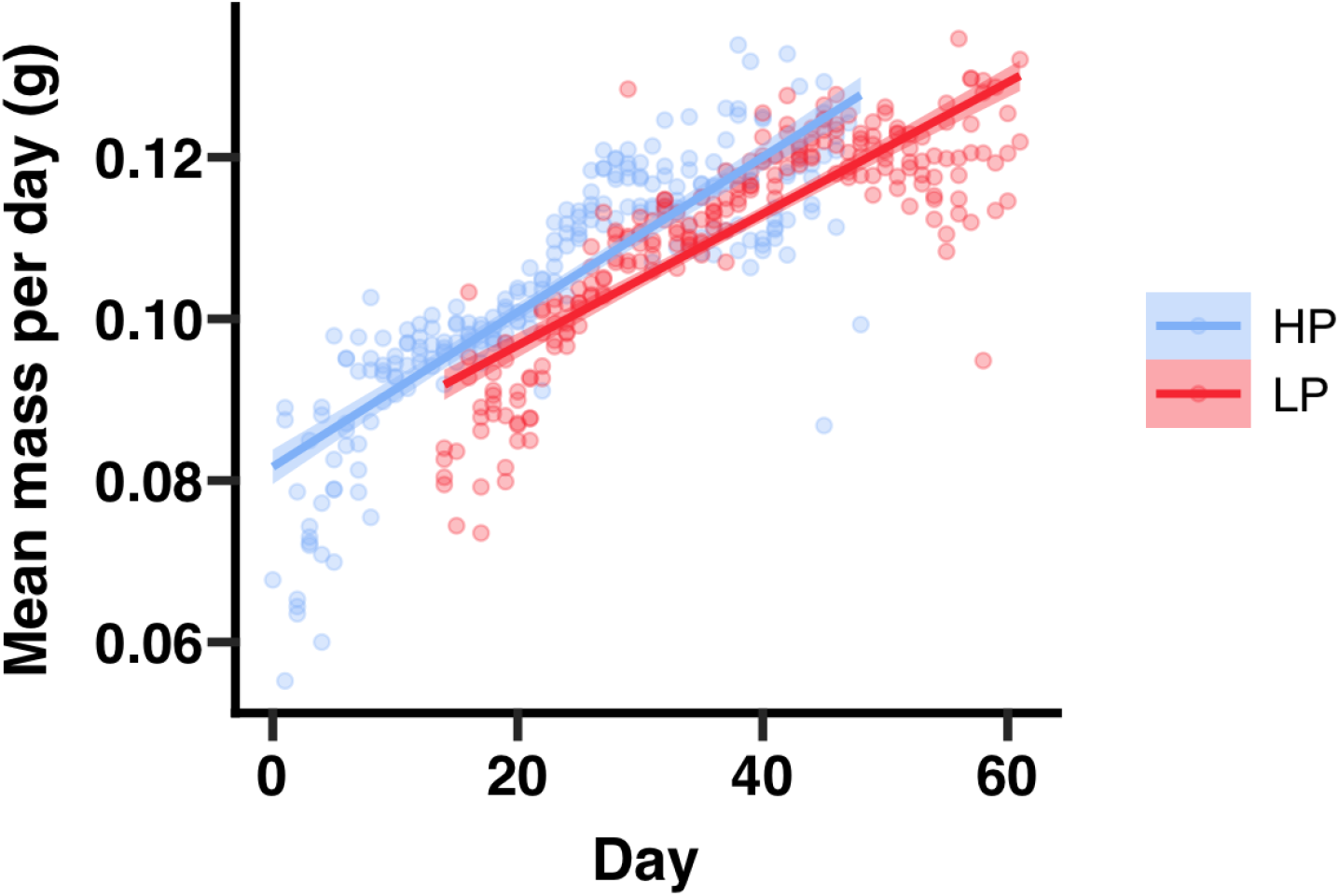
Average adult mass on emergence differs between treatments. **A)** Mean adult emergence mass increased over time (ANCOVA; t_445,449_ = 25.09, p < 0.0001).YML fed the HP diet showed a greater increase in mean mass on emergence with time than those fed the LP diet (ANCOVA; t_445,449_ = 2.68, p < 0.001). YML fed the LP diet had a higher mean adult body mass than those fed the HP diet (ANCOVA; t_445,449_ = 2.67, p < 0.001).

### Effect of larval and adult diet on oviposition

There was a significant difference in the numbers of offspring produced per female between HP- and LP- fed adult females, all of which were fed LP diet as larvae (LMM, F_1,30_ = 11.98, p < 0.01). Females fed the HP diet as adults had more offspring per female than females fed the LP diet in their first ten days of life (Figure 6A; t_24,30_ = 3.46, p < 0.01).

**Figure 6.**
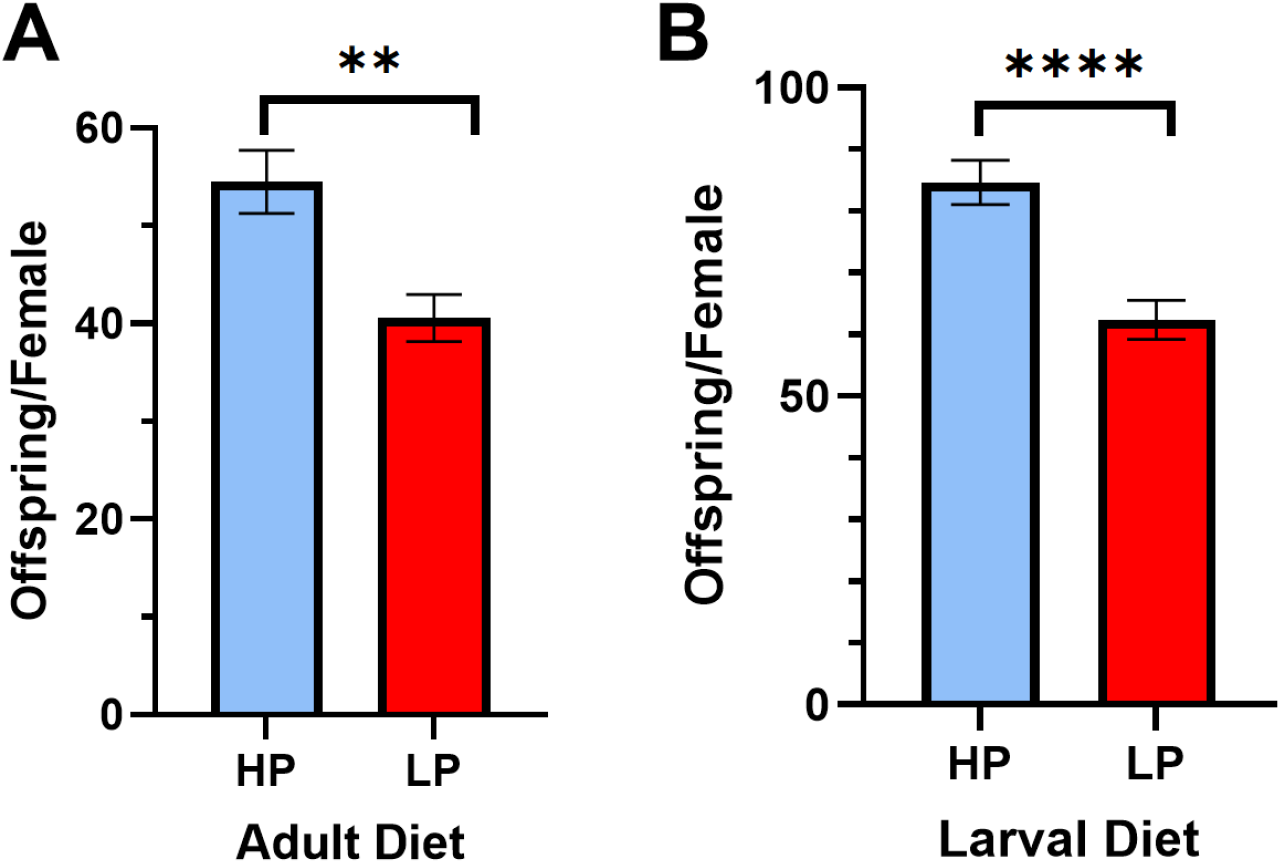
Females produce more offspring when fed HP diets either as adults (following a LP diet as larvae) or as larvae (when then fed a LP diet as adults). **A)** When LP-fed larvae were transitioned to a HP-fed diet as adults, they produced 54.5 ± 12.5 (mean ± SD) offspring per female (over 10 days), whereas adults fed the LP diet only produced 40.6 ± 9.27. There was a significant difference between the two treatments (LMM, F_1,30_ = 11.98, p < 0.01). **B)** Adults that were fed a HP diet as larvae produced 84.6 ± 13.9 offspring per female (over 15 days) compared to adults fed a LP diet as larvae, which produced only 62.4 ± 16.1 (all adults fed LP diet; LMM, F_1,29_ = 21.54, p < 0.0001). ** = p < 0.01, **** = p < 0.0001

There was a significant difference in the number of offspring per female between adults that had been fed a HP diet as larvae and adults that were fed a LP diet as larvae, when all were fed LP diets as adults (LMM, F_1,29_ = 21.54, p < 0.0001). Females fed the HP diet had more offspring per female than females fed the LP diet as larvae (Figure 6B; t_24,29_ = 4.63, p < 0.01).

## Discussion

Our data show that supplementing the YML diet with additional protein can have positive effects on growth rate, survival, and total larval yield at eight weeks. The higher protein diet reduced larval cannibalism rates and hastened development time. Larval cannibalism is reduced and survival is improved whether the HP diet is provided during earlier (weeks 0-4) or later (weeks 4-8) periods in larval development; though, providing supplemental protein earlier in life seems to be most consistently impactful for preventing cannibalism, improving survival, growth, and larval yield. If protein supplementation across the entire larval lifestage proves too costly, producers are likely to obtain the most benefit (in regard to survival, growth, and cannibalism reduction) from providing protein supplementation earlier during development. Corroborating our results on the significance of protein in the early-life diet for YML, a nutrient self-selection study found that they preferred a P:C ratio of close to 1:1 over the first 15 days of life and a ratio of 1:1.24 P:C over a more prolonged 30 day development period (Rho & Lee, 2022). These data indicate that prior welfare recommendations of at least 20% protein in the diet for YML (Barrett *et al*., 2023) may be insufficient, especially for earlier instars.

Importantly, while our data suggest cannibalism rates are reduced by protein-supplemented feed, they cannot show whether that cannibalism occurred prior to or after the death of the cannibalized individual. Therefore, it’s not clear if high protein diets drive reductions in cannibalism by reducing the ‘motivation to cannibalize’ (by providing access to other sources of protein to eat besides the body of living conspecifics) and/or by reducing mortality overall, and thereby reducing opportunistic but non-specific cannibalism of dead conspecifics. Although cannibalism of live individuals has been documented before for YML (reviewed in Barrett et al. 2024), cannibalism of dead or dying individuals appears more common (Barrett, pers. comm.); though, cannibalism of live individuals during/right after molts may also occur. In our study, we suspect reductions in cannibalism for YML on HP diets are driven by improved survival and thereby reduced opportunity to cannibalize, rather than vice versa. However, cannibalism of dead or dying conspecifics still represents a welfare issue insofar as it may hasten the spread of disease in a population (reviewed in Barrett et al. 2024) or add further to the potential suffering of dying individuals (Ichikawa & Kurauchi 2009, Weaver & McFarlane 1990). Therefore, reducing the prevalence of cannibalism is still expected to result in positive welfare impacts for YML.

One challenge with interpreting our data on adult body mass and pupation/eclosion timing among larval diets is a possible methodological limitation associated with the refeeding rates of YML, (Figure 1). HP larvae grew more rapidly and therefore, at later instars, likely consumed all their available feed more rapidly. We suspect there was a period of time between days 55 and 62 where the HP larvae had consumed all their available feed and the LP larvae had not, prior to a refeeding of all larval bins on day 63. For many larval insects, if they reach ‘critical mass’ (the minimum weight at which growth is no longer necessary to progress onward to pupation) and cannot find additional food, they will enter metamorphosis early (De Moed *et* al., 1999; Davidowitz *et al*., 2003), despite the capacity to grow larger under well-resourced conditions.

A seven day period of insufficient nutrition would explain the much earlier, first pupation timing for the HP larvae, which was not observed in our separate laboratory colony that regularly receives nutritional yeast supplementation; the bimodal peaks of adult emergence seen only for the HP condition (Figure 4; with the peaks representing, first, those that had and, second, those that had not, reached critical weight prior to the day 63 refeed); and the difference in observed individual biomass between our ‘small plate’ studies where food was always *ad libitum* due to the small number of individuals, and our ‘big bin’ studies. Therefore, we suspect that if larvae were provisioned based on ‘feed availability per condition’, instead of selecting standardized refeeding dates and amounts across both conditions, we may have seen later pupation timing, a single adult emergence peak, and larger adult body sizes in the HP condition compared to the LP condition. A future study could alter refeeding rates to determine how this impacts adult emergence timing and size more clearly when YML are fed with dietary protein supplements. These data are still valuable for producers, however, demonstrating that it is possible to produce faster pupation timing by feeding HP diets early and then reducing feed quantity later after larvae have hit critical mass, thereby replicating the conditions in this study.

We also found that providing a HP diet as either larvae, or for adults (even if they have previously been reared on a LP, standard diet), can increase egg yield by 34-36% (and see: Morales-Ramos *et al*., 2013). These data are unlikely to be due to differences in egg cannibalism by adults, due to the use of a mesh excluder; therefore, this represents true differences in fecundity over the first 10 or 15 days of a female’s lifespan. Other studies confirm that protein is especially important for female insects to be able to lay eggs, and is often selected for at higher rates by female insects compared to males in adult nutritional choice assays (Maklakov *et al*., 2008; Camus *et al*., 2018; Sydney *et al*., 2024), including yellow mealworm adults (Rho & Lee, 2016). Our data additionally show that larval dietary protein supplementation can improve adult fecundity even when supplemental protein is excluded from adult diets.

These data suggest that industry-standard diets of pure wheat bran may not provide enough protein for YML, driving reductions in growth, survival, and yield and increasing welfare-negative behaviors like cannibalism. Previous studies have also found positive effects of protein supplementation for YML, for both production and putative welfare reasons (Morales-Ramos *et al*. 2013, Mahmoud *et al*., 2025, Rho & Lee, 2022). We demonstrate that these effects are most critical during the first four weeks of life, but there are also benefits when protein supplementation is provided later in larval development. Further, we highlight that providing protein supplements to yellow mealworm adults - in line with nutrient self-selection studies (Rho & Lee, 2016) - results in increased fecundity, demonstrating the benefits of greater protein availability across the life cycle. Further research should focus on nutrient self-selection by larvae at different instars, as well as assess the degree to which living/dying, compared with dead, individuals are subject to cannibalism and thus the magnitude of the welfare risk. The precise timing and concentration of protein, as well as a better understanding of micronutrients that may play a role in supporting YM development and occur concomitantly in some protein sources like yeast, should also be investigated to ascertain the most cost-effective application of dietary protein supplementation for YML.

## Acknowledgments

Thanks to Mariangeles Paloma Zacarias and Elijah Persson-Gordon for assistance counting larvae, especially on some weekends, for the cannibalism experiment.

